## Supplementary table 1 for "TRPML1 gating modulation by allosteric mutations and lipids"

|  | **TRPML1**  **Y404W** | **PI(4,5)P_2_-bound**  **TRPML1** |
| --- | --- | --- |
|  | EMDB-45429  PDB-9CBZ | EMDB-45432  PDB-9CC2 |
| **Ligand treatment condition** | None | 0.5mM PI(4,5)P_2_ diC8 |
| **Data collection and processing** |  |  |
| Magnification | 105,000 | 165,000 |
| Voltage (kV) | 300 | 300 |
| Electron exposure (e–/Å2 ) | 60 | 60 |
| Defocus range (μm) | -0.9 - -2.2 | -0.9 - -2.2 |
| Pixel size (Å) | 0.826 | 0.738 |
| Symmetry imposed | C4 | C4 |
| Initial particle images (no.) | 864,698 | 1,065,778 |
| Final particle images (no.) | 35,460 | 60,597 |
| Map resolution (Å) | 2.86 | 2.46 |
| FSC threshold | 0.143 | 0.143 |
| **Refinement** |  |  |
| Initial model used (PDB code) | 5WPV | 5WPV |
| Model resolution (Å)  FSC threshold | 2.86 | 2.46 |
| Map sharpening B factor (Å2 ) | -82.51 | -60.00 |
| Model composition |  |  |
| Non-hydrogen atoms | 15,424 | 15,556 |
| Protein residues | 1,892 | 1,856 |
| Ligands | 8 | 16 |
| B factors (Å2 ) |  |  |
| Protein | 26.70 | 30.90 |
| Ligand | 40.90 | 77.59 |
| R.m.s. deviations |  |  |
| Bond lengths (Å) | 0.005 | 0.002 |
| Bond angles (°) | 0.633 | 0.457 |
| Validation |  |  |
| MolProbity score | 1.47 | 1.24 |
| Clashscore | 5.89 | 4.57 |
| Poor rotamers (%) | 0 | 0 |
| Ramachandran plot |  |  |
| Favored (%) | 97.23 | 97.98 |
| Allowed (%) | 2.77 | 2.02 |
| Disallowed (%) | 0 | 0 |

Table S1. Data collection and refinement statistics.
